# The role of anxiety in the perception of pain: exploring the cumulative & temporal mechanisms of hypercapnic analgesia

**DOI:** 10.1101/2020.10.30.357061

**Authors:** Anna Yurievna Kharko, Kirralise J Hansford, Frederico B Klein, Paul L Furlong, Stephen D Hall, Matthew E Roser

## Abstract

**Background:** Anxiety, evoked by continuous inspiration of a 5 – 8% CO_2_ mixture, has been found to have an analgesic eLect on self-reported pain. The precise mechanism whereby this effect obtains remains unknown.

**Methods:** The present study tested whether temporal summation, the psychological counterpart of wind-up, is involved in hypercapnic analgesia. 21 healthy participants received painful transcutaneous electrical stimuli of varied intensity, during continuous inhalation of 7.5% CO_2_ mixture and medical air, presented in a single-blinded counterbalanced order. Continuous pain ratings were acquired to measure the temporal development of the pain response. Several points and events of interest that characterise the pain response profile were extracted from the continuous data.

**Results:** Mixed-eLects modelling demonstrated a reduction of all pain measures during inspiration of the anxiogenic mixture, but not air. This was accompanied by an increase in the psychological and physiological measures of anxiety. Analyses of the characteristic measures of temporal summation suggested that the hypercapnic mixture has an analgesic property evident from the start of the pain response. The same was true for the remainder of the response, the adaptation period, where pain ratings were also inhibited. The reduced pain ratings persisted during the remainder of the response. Anxiety was found to be a mediating factor for summative pain ratings but not the temporally sensitive TS measures, suggesting an overall, cumulative effect.

**Conclusions:** The findings provide an explanation for the previously observed low self-reported pain during the inhalation of an anxiogenic hypercapnic mixture.

## Introduction

In psychopharmacology, the 5 - 10% carbon dioxide (CO_2_) model is an established human experimental model of anxiety (Bailey et al., 2011). It requires the participant to continuously breathe in the CO_2_ mixture, which elicits a range of stable physiological symptoms of anxiety including hypercapnia, hypertension, tachycardia and diaphoresis (Pappens et al., 2012; Poma et al., 2005; Tominaga et al., 1976; Woods et al., 1988). Psychological and behavioural markers of anxiety are also observed, including increased ratings and attribution of negative affect, poor emotion recognition, and impaired information processing (Attwood et al., 2017; Cooper et al., 2013; Easey et al., 2018).

In pain research, as early as 1948, CO_2_-evoked hypercapnia has been found to elevate pain threshold (Stokes et al., 1948). More recently, studies have demonstrated a dose-dependent decrease in self-reported pain intensity during the administration of thermal (Grönroos and Pertovaara, 1994) and mechanical pain (Vowles et al., 2006). While CO_2_ inspiration appears to have an analgesic eLect, electrophysiologic data does not definitively support this. The inhalation of a hypercapnic mixture has been found to produces an anti-nociceptive eLect characterised by preserved nociceptive flexion reflex (Morélot-Panzini et al., 2014).

The decrease in self-reported pain, despite the persistent nociceptive reflex, has suggested that CO_2_-mediated analgesia is preserved at a supraspinal level (Grönroos and Pertovaara, 1994); however, this has not yet been explicitly tested. One way to do so is by examining temporal summation (TS); the escalation of consecutive pain ratings in response to continuous stimulation of constant intensity at a critical frequency, within the first seconds of the painful input (Price et al., 1977). TS is the behavioural counterpart of wind-up; a central nervous system (CNS) measured increase in excitability of dorsal-horn neurons during nociceptive stimulation. While the CO_2_ model has been used to study wind-up, it has not yet been applied to TS.

Further, since TS originates from self-report, is likely to be affected by the subjective experience of the hypercapnic inhalation. Previous research has found a complex aversive response to pain and/or CO_2_-induced fear (Vowles et al., 2006).

However, the anxiogenic effects of the inhalation and the temporal development of acute pain concurrent to CO_2_-induced anxiety have not been directly investigated.

Here, we address the questions using a robust continuous pain-measurement approach (Wijk et al., 2013), to capture the influence of the hypercapnic mixture on TS of pain and self-reported acute anxiety. We recorded the moment-to-moment changes in continuous pain ratings of painful transcutaneous electrical stimulation (TES), during inspiration of a 7.5% CO_2_ mixture (anxiogenic condition) and medical air (control condition).

Consistent with previous reports, we hypothesised that summative pain ratings would be lower in the anxiogenic than the control inhalation. Utilising the continuous pain data, we next hypothesised that key measures extracted from the TS period would also reveal an analgesic effect of the CO_2_ inhalation. In particular, TS rate would be slower (reduced TS slope) and reaching a lower maximum (reduced TS peak). Further, the pain ratings in the period following TS will remain reduced in the CO_2_ condition. Lastly, we predicted that the lowered pain measurements will be associated with heightened anxiety.

## Methods

### Participants & screening

An extensive screening process was completed to minimise the risk of adverse reaction to the experimental process. Consequently, the participant group satisfied the following criteria: participants were 18 to 60 years of age; with a healthy BMI (18 – 30); were not pregnant (confirmed or suspected) or breastfeeding. Participants had no diagnosis of a concurrent chronic or acute condition; no medical issues relating to respiratory or cardiovascular systems; had no Axis I or Axis II diagnoses; and reported no acute pain at the time of testing. Participants were non-smokers (including vaping); had no current or history of substance abuse, including illicit drugs intake, alcohol abuse (> 35 units/week for females), and no excess caLeine consumption (> 8 cups/day). Participants had taken no medication in the past month, other than contraceptive pills, topical creams or incidental use of over the counter mild analgesics and antihistamines. In compliance with the guidelines set by the manufacturer of the stimulation equipment, participants had no pacemakers, shunts, stents or metal in the body, and no skin abrasions on the site of TES.

In preparation to testing, participants were required to discontinue alcohol consumption 24 hrs prior to the study and xanthine-based beverages on the day of the study. Final eligibility was confirmed using the Mini-International Neuropsychiatric Interview (MINI) (Sheehan et al., 1998). Blood pressure (BP) and heart rate (HR) were monitored throughout the study, to ensure they remained within an acceptable range, as determined by the UK National Health Service (NHS) guidelines (BP range 90/60 - 140/90 mmHg; HR 60 - 100 bpm). Variance outside of these ranges would result in termination of the experiment.

Following screening, the participant group consisted of twenty-six adult females. Participant data, including age, ethnicity, height, weight, and BMI were recorded for further analysis. Participants were invited to a single testing session. Collected data was anonymised. Ethical approval (Ref. Number 18/19–1044) was obtained from the School of Psychology, Faculty of Health Ethics Committee at the University of Plymouth, UK. Informed written consent was obtained on the day of testing.

### Pain history

Data from this study will also be used to inform the development of a further implementation of the CO_2_ model in clinical pain populations. Therefore, additional data related to fibromyalgia (FM) were recorded but not reported here. These included the American College of Rheumatology Criteria from 1990, ACR’90 (Wolfe et al., 1990) and from 2010, ACR’10 (Wolfe et al., 2010), considering the redactions from 2016 (Wolfe et al., 2016). Recent pain history was also assessed through the Short-Form McGill Pain Questionnaire, SF-MPQ (Melzack, 1987).

### TES stimulation & pain rating

All psychophysical testing was performed by the same experimenter. Throughout the process participants were seated in a reclining chair in a semi-Fowler’s position.

#### Stimulation site

Stimulation site was the skin over the sural nerve at lateral border of the tendo-Achilles. Body side was counterbalanced between participants and stimulation site was confirmed to be free from hair and skin trauma. The site was prepared with an abrasive gel and cleaned with isopropyl alcohol prior to electrode placement. Two disposable self-adhesive disk electrodes, 2 cm in diameter, were positioned with inter-electrode distance of 3 cm. Electrical stimulation was delivered by a constant current stimulator DS7A Digitimer (Digitimer Ltd., UK). During acquisition of psychophysical measures, the equipment was manually operated by the experimenter.

#### Psychophysical measures

Three psychophysical measures were collected to characterise individual pain profiles and calculate stimulation levels: *sensory threshold* (STHR), *pain threshold* (PTHR), and *maximum tolerance* (MTOL). STHR was the current amplitude at which the participant first felt a sensation at the site of stimulation. PTHR was the amplitude at which the participant perceived the stimulation as minimally painful. MTOL was the amplitude at which the participant was unwilling to experience a further stimulation intensity increase. *Pain range* (PRAN) was then determined as the difference between MTOL and PTHR (PRAN = MTOL – PTHR). Participants were asked to describe the sensation at each measure to ensure correct interpretation of instructions. Current amplitudes were identified using a staircasing approach, with pulses delivered at 1Hz, with increments of 0.1mA. A maximal acceptable current of 40mA was pre-defined for safety. Psychophysical measures were acquired three times and their mean value was used to determine the stimulation levels used during the pain rating task.

#### Stimulation parameters

There were six experimental stimulation levels (SLs) in ten-percent increments (10-60%), calculated as:

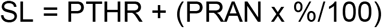

Experimental stimuli were single square-wave pulses with a pulse-width of 500μs, delivered at 2.5Hz. Stimuli were delivered at each of the stimulation levels in 45-second blocks, in a pseudo-randomised order. Each condition was repeated twice, producing a total of 12 experimental blocks. Stimulation was delivered semi-automatically using a computer, with manual adjustment of intensity between blocks.

#### Pain ratings

Once the psychophysical measures and stimulation levels were determined, the participant was introduced to the pain rating task. A trial run of two blocks was carried out without anxiety induction.

A pain visual analogue scale (pVAS) was used to continuously measure pain ratings on a scale of 0 to 100, delineated with ticks at tenth intervals, where ‘0’ represented lack of pain, ‘1’ was the minimal sensation of pain, and ‘100’ was the worst pain imaginable. The rating was provided using a custom-build response box fitted with a rotary potentiometer button. Participants continuously rated the stimulation for the duration of the block, with values being recorded every 50 ms using an Arduino Genuino (Arduino LLC).

### CO_2_ model & anxiety testing

#### Gas mixtures

Two gas mixtures were used in the experiment. The anxiogenic mixture contained elevated CO_2_ (7.5% CO_2_/21% O_2_/ 71.5% N_2_). The control mixture was Medical Air (0.04% CO_2_/21% O_2_/ 78% N_2_). Gas cylinders were connected to a 10 L Douglas Bag and delivered to participants via a sterilised reusable face mask (7450 Series V2, Hans Rudolph Ltd). Inspiratory and expiratory flows were isolated using two-way non-rebreathing valves (Hans Rudolph Ltd). Delivery order of gas mixtures was randomised, single-blinded for safety purposes and counterbalanced between participants.

#### Anxiety measures

Several measures of anxiety were collected. Participants completed the Trait version of the State-Trait Inventory for Cognitive and Somatic Anxiety, STICSA-T (Grös et al., 2007), prior to psychophysical testing to measure dispositional anxiety. Changes in state anxiety were measured throughout the study, using the State version of STICSA, STICSA-S (Grös et al., 2007), BP, HR, as well as an anxiety VAS (aVAS). aVAS was also a 101-point scale ranging from ‘0’ – no anxiety, through ‘1’ – least anxiety, to ‘100’ – worst anxiety imaginable. Anxiety measures were delivered at key points during the study. STICSA-S was administered five times: prior to psychophysical testing, immediately following each inhalation session, and following each post-inhalation break. BP and HR were also recorded at five intervals: prior to psychophysical testing, two-minutes into each inhalation, and following each post-inhalation break. Twenty-seven individual aVAS ratings were recorded: prior to psychophysical testing and during each inhalation at the start of each stimulation block (12 blocks during each inhalation or 24 blocks in total), as well as at the end of each post-inhalation break.

### Procedure

Participants first completed pain history and anxiety scales, followed by psychophysical testing and a pain rating practice run. After familiarisation with the task and safety briefing, participants were connected to the respiratory equipment. An initial two-minute idle inhalation, followed by BP and HR measurement, was used to confirm that participants’ cardiovascular measures were within the acceptable range. Eligible participants continued to the testing session, which comprised of the first up to 15-min gas inhalation (CO_2_ mixture or Medical Air), followed by a mandatory 20-min break, then second up to 15-min gas inhalation (CO_2_ mixture or Medical Air), then final mandatory break. Participants were debriefed and reimbursed upon study termination. The experimental procedure is visualised in Figure 1.

**Figure 1.**
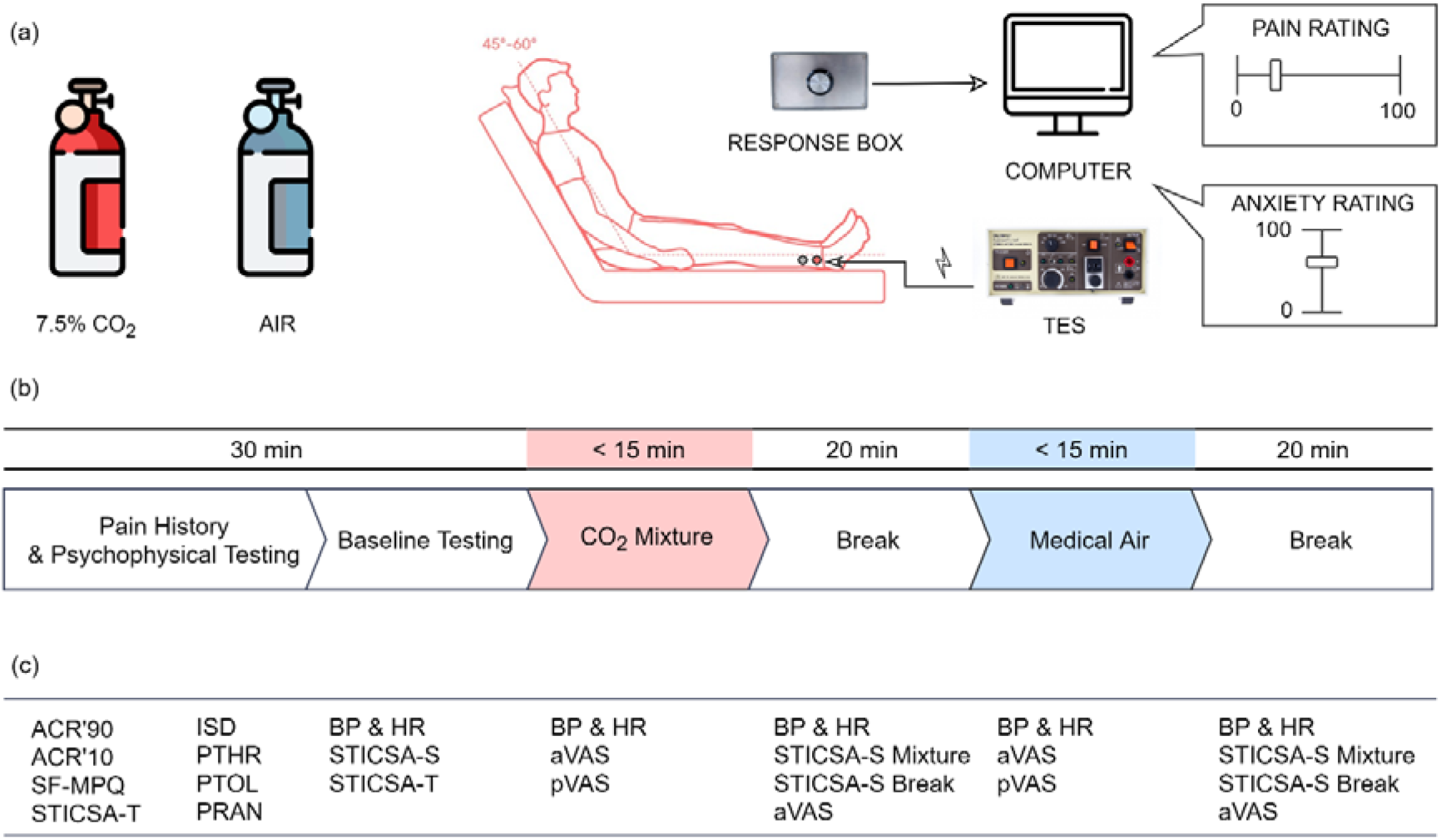
(a) The equipment setup. (b) The study stages. (c) Acquired measures. TES: transcutaneous electrical stimulator; ACR’90/’10: American College of Rheumatology Questionnaire from 1990/2010; SF-MPQ: Short Form McGill Pain Questionnaire; STICSA-S/T: State-Trait Inventory for Cognitive and Somatic Anxiety - State / Trait Version; STHR: sensory threshold; PTHR: pain threshold; MTOL: maximum tolerance; PRAN: pain range; BP: blood pressure; HR: heartrate; a/pVAS: anxiety/pain visual analogue scale. Stimulator image is supplied by manufacturer (Digitimer Ltd., UK), cylinder icons are sourced from FlatIcon, and sitting position image from Dimensions.Guide. Reproduced from respective sources with permission.

### Analysis

All analysis was carried out using RStudio v1.0.153 (RStudio Inc). Descriptive statistics for the sample, questionnaire scores, psychophysical measures, and stimulation levels were summarised as: mean values (μ) or count (n) and standard deviation (*SD*) or percentage (%); as appropriate. Mixed-effects modelling was used to determine the factors that affected a given outcome variable. Each model contained two fixed effects - *Mixture* (CO_2_ Mixture or Medical Air), and *Stimulation* (SL10%, 20%, 30%, 40%, 50% or 60%). There were several possible random effects: *Participant*, and the anxiety measures *aVAS*, *STICSA-S*, *HR*, systolic BP (*SYS)*, and diastolic BP (*DIA)*. Model selection was theory-driven (Barr et al., 2013), toward a parsimonious formula (Bates et al., 2018), that minimises Type I error in small samples (Brysbaert and Stevens, 2018; Matuschek et al., 2017). The observed power for each appropriate model was assessed using Kenward-Roger approximation. Hypothesis testing was achieved by comparison to a null model (by-participant intercept only), based on Bayes Factor (BF), likelihood-ratio testing (χ2), and p-values. The final model statistics were: estimated μ and coefficient, SD and standard error (SE), reported alongside Bayesian estimated μ and coefficient, SD, and credible intervals. Bayesian analysis was carried out using weakly informed priors (Muth et al., 2018).

#### Average pain ratings

In our first hypothesis we predicted that summative pain ratings would be lower in the anxiogenic than the control inhalation. This was tested by comparison of the mean pain ratings per block (pVASμ) and per period. The continuous pain response was broken into two periods. At the start of a response block we expected to observe a period of rapid incline of pain ratings. We termed this portion of the pain response the TS Period. Taking into account the stimulation protocol (Arendt-Nielsen et al., 2000), the length of the TS Period was predetermined to span the initial 15 s. A measure of mean pain rating during the TS period (TSμ) was calculated alongside with the averaged pVAS rating for the time interval between 15 s - 45 ms called a period of adaptation, or A Period (Aμ). A prototypical response with the described periods could be seen in Figure 2.

**Figure 2.**
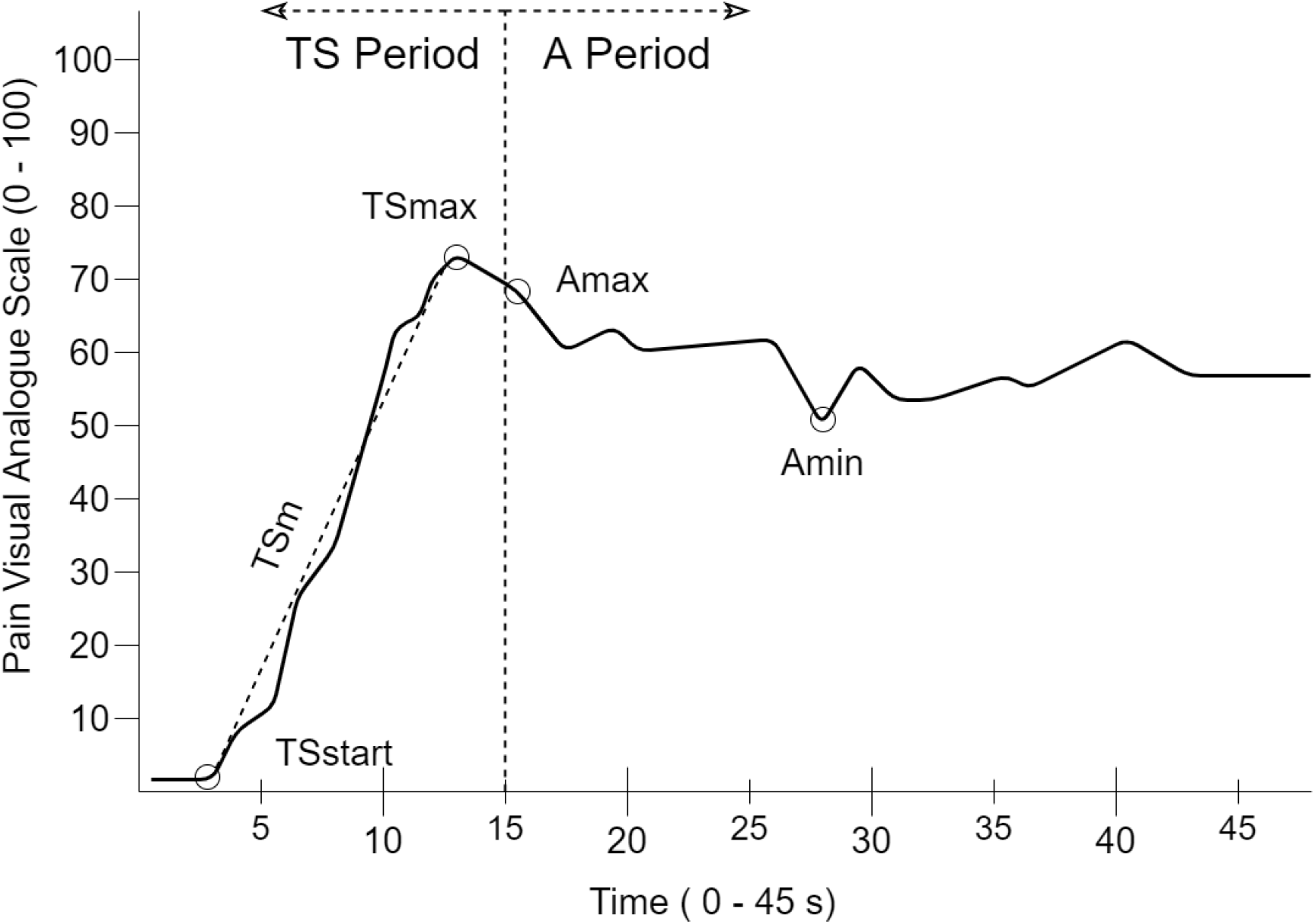
The prototypical continuous pain rating response is broken into two periods with several measures of interest. TS Period: temporal summation period; A Period: adaptation period; TSstart: the start of the response; TSmax: the maximal rating made during TS; TSm: the slope calculated between the start of the response and the maximal TS value; Amin: the minimal value; Amax: the maximal value were recorded.

#### Continuous pain ratings

Our second hypothesis predicted that key measures extracted from the TS Period will be reduced during the CO_2_ inhalation. This was tested through comparison of the peak and the slope of the pVAS during TS. The peak (TSmax) was the maximal pVAS rating during TS, and the slope (TSm) was computed by fitting a line between the first pain rating (TSstart) to TSmax.

In our third hypothesis we predicted that the CO_2_ mixture will continue to lead to reduced pain ratings in the A Period. Here we computed two measures to capture the pVAS response: maximal (Amax) and minimal (Amin) pVAS ratings.

#### Anxiety measures

In the last hypothesis we stated that increased anxiety will be associated lowered pain measurements. To test this we first confirmed that the CO_2_ model was successful, by analysing whether the CO_2_ inhalation significantly increased the psychological (aVAS, STICSA-S) and physiological (HR, DIA, SYS) measures of acute anxiety. This was determined using separate mixed-effects models. The inclusion of fixed and random effects following the same approach as previously described, with the addition of pVASμ as a random effect to account for the potential interaction of pain and anxiety. To test whether the elevated acute anxiety is associated with reduced pain measurements, the above mentioned anxiety measures were included as potential random effects as previously described.

Data relevant to the hypotheses are stored on Open Science Framework: https://osf.io/ds4e3/?view_only=07d2a59bd8a54685ac44700bc910947d.

## Results

### Participants data, pain history & psychophysical testing

Five of the 26 participants did not complete the 7.5% CO_2_ inhalation, providing a final sample of 21 females. Their demographic data, cumulative scores on pain history questionnaires and average psychophysical measures are summarised in Table 1.

**Table 1.** Descriptive statistics for demographics, pain history, and psychophysical measures.

| | $\mu$ or n | (SD) or % |
| --- | --- | --- |
| Age (n years) | 25.24 | (7.27) |
| Ethnicity <sup>a</sup> |  |  |
| Asian British | 1 | 5% |
| Black British | 1 | 5% |
| White British | 18 | 85% |
| White Other | 1 | 5% |
| STICSA-T | 26.76 | (5.52) |
| Cognitive Anxiety | 13.33 | (3.45) |
| Somatic Anxiety | 13.43 | (2.6) |
| SF-MPQ |  |  |
| Pain Descriptors (0 - 45) | 2.14 | (2.15) |
| Visual Analogue Scale (0 - 10) | 1.05 | (.93) |
| Present Pain Intensity (0 - 5) | .1 | (.3) |
| STHR (mA) | 1.30 | (.42) |
| PTHR (mA) | 6.56 | (3.57) |
| MTOL (mA) | 14.62 | (7.57) |
| PRAN (mA) | 8.06 | (6.83) |

Note: $\mu$ - average value, n – count, SD – standard deviation, % - percentage.
<sup>a</sup> Items, for which n and % were calculated.

### Pain Ratings

#### Average pain ratings

The best mixed-effects model for each average pain measure had the same fixed and random effects but differed in performance over baseline model (see Table 2).

**Table 2.** Formulas and Statistics for Mixed-effects Models for Pain Measures

| Variable | Model Formula | $\chi^2$ | $p$ | BF |
| --- | --- | --- | --- | --- |
| **Average Pain Ratings** |  |  |  |  |
| pVASμ | ~ Mixture + Stimulation + (1 \| Participant/aVAS) | 254.28 | < .001 | $1.68 \times 10^{50}$ |
| TSμ | ~ Mixture + Stimulation + (1 \| Participant/aVAS) | 227.74 | < .001 | $8.42 \times 10^{43}$ |
| Aμ | ~ Mixture + Stimulation + (1 \| Participant/aVAS) | 249.19 | < .001 | $2.99 \times 10^{49}$ |
| **Continuous Pain Ratings** |  |  |  |  |
| TSmax | ~ Mixture + Stimulation + (1 \| Participant) | 234.49 | < .001 | $6.48 \times 10^{42}$ |
| TSm | ~ Mixture + Stimulation + (1 \| Participant) | 19.48 | < .001 | .01 |
| Amax | ~ Mixture + Stimulation + (1 \| Participant) | 250.45 | < .001 | $1.9 \times 10^{46}$ |
| Amin | ~ Mixture + Stimulation + (1 \| Participant/aVAS) | 249.19 | < .001 | $3 \times 10^{49}$ |

Note: pVASμ : average pain rating for the whole response; TSμ : average pain rating for the period of temporal summation; Aμ : average pain rating for the period of adaptation. The choice of best model was design-driven with the goal of reaching a formula with a parsimonious number of factors. Each model was compared to a baseline model based on likelihood-ratio testing ( $\chi^2$ ) and BIC-derived Bayes Factor (BF).

The best model for pVASμ found that it decreased during the CO_2_ Mixture (*Coeff.* = −3.68, *SE* = 1.23, *t* = 3.01, *p* = .003) but significantly increased with each Stimulation level. Variance in the pVAS was explained by individual variability, Participant (σ^2^ = 180.5, *SD* = 13.43), and individual anxiety rating, aVAS (σ^2^ = 55.4, *SD* = 7.44). pVASμ ratings per stimulation level and inhalation are summarised in Table 3.

**Table 3.** Average stimulation levels and pain ratings per inhalation.

| Stimulation |  |  | CO<sub>2</sub> Mixture |  | Medical Air |  |
| --- | --- | --- | --- | --- | --- | --- |
| | $\mu$ | (SD) | $\mu$ | (SD) | $\mu$ | (SD) |
| SL 10% | 7.36 | (3.61) | 16.59 | (13.86) | 18.87 | (16.52) |
| SL 20% | 8.17 | (3.77) | 22.27 | (15.73) | 26.14 | (17.95) |
| SL 30% | 8.97 | (4.04) | 28.29 | (18.41) | 35.36 | (19.29) |
| SL 40% | 9.78 | (4.40) | 37.35 | (20.8) | 39.51 | (18.44) |
| SL 50% | 10.59 | (4.83) | 40.14 | (18.15) | 41.13 | (20.39) |
| SL 60% | 11.39 | (5.32) | 44.63 | (21.34) | 52.14 | (22.46) |

Note: SL: stimulation level, calculated as a percentage of an individual's pain range added to the individual's pain threshold. Descriptive statistics include $\mu$ : average value, and SD: standard deviation.

The same was true for TSμ and Aμ. TSμ was reduced during the CO_2_ Mixture (*Coeff.* = −2.9, *SE* = 1.07, *t* = 2.72, *p* = .007) but significantly increased with each increase of Stimulation intensity. Participants (σ^2^ = 106.39, *SD* = 10.31) and their anxiety rating, aVAS (σ^2^ = 64.06, *SD* = 8), accounted for large proportion of the variance in TSμ. Similarly, the CO_2_ Mixture led to a decrease in Aμ (*Coeff.* = −4.11, *SE* = 1.36, *t* = 3.02, *p* < .01) while increase of Stimulation lead to an increase in Aμ, when considering the variance explained by Participants (σ*2* = 236.63, *SD* = 15.38) and their anxiety rating, aVAS (σ^2^ = 51.64, *SD* = 7.19). Full model statistics can be viewed online in Supplemental Material 1.

#### Continuous pain ratings

The best mixed-effects models for continuous pain data differed in random effects (see Table 2). Detailed statistics for each model can be found online in Supplemental Material 2.

Mixed-effect modelling found that TSmax was lower during the inhalation of the CO_2_ Mixture (*Coeff.* = −4.56, *SE* = 1.21, *t* = 3.78, *p* = .001), increased with each Stimulation increase, with large portion of the rating explained by individual variability, Participant (σ*2* = 217.4, *SD* = 14.74). aVAS was not found to significantly improve the model unlike it did in the summative pain measures analysis. TSm was also decreased during the CO_2_ Mixture (*Coeff.* = −1.37, *SE* = .51, *t* = 2.71, *p* = .007), but significantly increased in the stronger intensity stimulation conditions SL 40%, SL 50%, SL 60% compared to the reference category SL10%, when accounting for Participant variability (σ^*2*^ = 4.12, *SD* = 2.03). As with TSmax, the model for TSm that included anxiety was not superior.

Analysis of the A Period found similar effects of the anxiogenic manipulation and the painful stimulation. The CO_2_ Mixture lowered Amax (*Coeff.* = −3.92, *SE* = 1.28, *t* = 3.06, *p* = .002) while all Stimulation levels significantly increased it. Participant variability further explained Amax (σ*2* = 234.6, *SD* = 15.32). For Amin, the CO_2_ Mixture also further reduced the rating (*Coeff.* = −4.11, *SE* = 1.36, *t* = 3.02, *p* = .003), while Stimulation had an increasing effect. Unlike Amax, Participant (σ^2^ = 236.63, *SD* = 15.38) was not the only random effect accounting for the variance in Amin. The individually reported anxiety, aVAS was also contributing to the rating (σ^2^ = 51.64, *SD* = 7.19).

### Anxiety testing

Both psychological measures (aVAS, STICSA-S) and physiological markers (HR, DIA, SYS) of acute anxiety were significantly increased during the inspiration of the CO_2_ mixture, compare to that of medical air. The variability of STICSA-S and cardiovascular values through the progression of the experiment can be seen in Figure 3.

**Figure 3.**
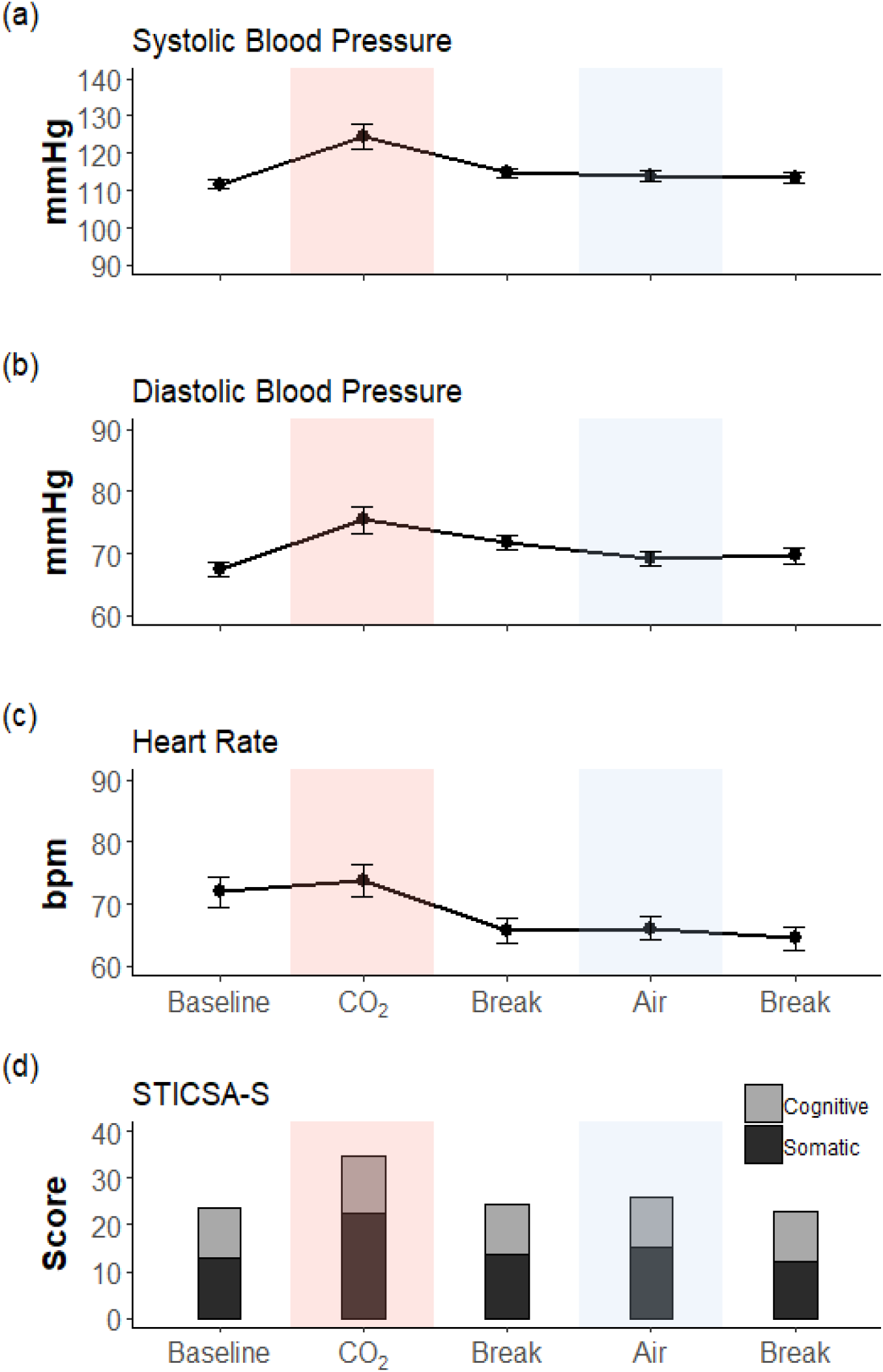
(a) On the average systolic blood pressure plot the y-axis shows the acceptable range from 90 to 140 mmHg. (b) On the average diastolic blood pressure plot the y-axis shows the acceptable range from 60 to 90 mmHg. (c) On the average heart rate plot the y-axis shows acceptable range. (d) On the y axis is the total score for STICSA-S: State-Trait Inventory for Cognitive and Somatic Anxiety – State Version. The total score is composed of the cognitive and somatic components of the inventory.

For all subsections, on the x-axis are plotted the study stages: Baseline: beginning of the study; CO_2_: the CO_2_ Inhalation; Break: the break after the CO_2_ Inhalation/Air; Air: the Air Inhalation. Error bars are standard error.

Mixed-effects modelling found that all of the best models for anxiety measures shared the same formula: ~ Mixture + (1 | Participant). The best models always included Mixture as a fixed effect, Participant as a random effect and never Stimulation as a fixed effect. The latter was due to Stimulation never being a significant predictor for any of the anxiety outcome variables. pVASμ, which was an additional optional random effect for these analyses, was never found to explain variance in of the anxiety measures.

The best model for aVAS found that the CO_2_ Mixture significantly increased the rating (*Coeff.* = 16.26, *SE* = .94, *t* = 17.33, *p* < .001) when taking into account Participants (σ^2^ = 124.8, *SD* = 11.17), Intercept (*Coeff.* = 8.75, *SE* = 2.53, *t* = .346, *p* = .002), χ2 = 233.98, *p* < .001, BF = 4.13 × 10^50^. The same was true for STICSA-S which increased during the CO_2_ Mixture (*Coeff.* = 8.62, *SE* = 0.20, *t* = 42.82, *p* < .001), after taking into account variance explained by Participants (σ^2^ = 28.80, *SD* = 5.37), Intercept (*Coeff.* = 25.95, *SE* = 1.18, *t* = 22, *p* < .001), χ2 = 757.26, *p* < .001, BF = 1.22 × 10^163^.

The cardiovascular measures followed the same pattern of increase. SYS was significantly higher during the CO_2_ Mixture (*Coeff.* = 16.14, *SE* = 0.46, *t* = 35.32, *p* < .001), when accounting for participants’ variance (σ^2^ = 51.75, *SD* = 7.19), Intercept (*Coeff.* = 113.91, *SE* = 1.6, *t* = 71.1, *p* < .001), χ2 = 616.41, *p* < .001, BF = 3.17 × 10^132^. As was DIA in the CO_2_ Mixture (*Coeff.* = 10.91, *SE* = 0.37, *t* = 29.74, *p* < .001), after accounting for Participants variance (σ^2^ = 23.12, *SD* = 4.81), Intercept (*Coeff.* = 69.24, *SE* = 1.08, *t* = 64.1, *p* < .001), χ2 = 502.63, *p* < .001, BF = 6.22 × 10^107^; and HR in the CO_2_ Mixture (*Coeff.* = 11.43, *SE* = .39, *t* = 29, *p* < .001), Participants’ variance (σ^2^ = 80.95, SD = 9), Intercept (*Coeff.* = 66.14, *SE* = 1.98, *t* = 33.4, *p* < .001), χ2 = 487.08, *p* < .001, BF = 2.61 × 10^104^.

## Discussion

The present study investigated the effects of an established CO_2_ experimental anxiety model on self-reported pain ratings using TES. We confirmed that increased state anxiety was elicited by the model, as previously reported. The addition of continuous pain ratings, using the pVAS approach, enabled high temporal-resolution characterisation of the emerging pain response alongside measures of anxiety. Analysis of continuous measures compared to cumulative measures revealed inconsistent effects of CO_2_-induced anxiety.

Consistent with previous observations, analysis of cumulative measures found that average pain ratings per block (pVASμ) are reduced in the anxiogenic condition compared to control, but increase with stimulation intensity. The same was true for the average ratings per TS Period and A Period (TSμ and Aμ). Mixed-effects modelling confirmed that the best model for prediction of all three averaged pain ratings included the aVAS, but not the anxiety questionnaire (STICSA-S), nor the cardiovascular values (SYS BP, DIA BP, HR). It is reasonable to suggest that this reflects a greater level of accuracy in the aVAS compared to other measures. Practical constraints on the timely completion of STICSA-S during the experiment likely had an impact on the utility of the resulting scores, as a consequence of changing conditions (during or after inhalation) and memory bias (Mitte, 2008). The use of aVAS enabled measurement of acute anxiety at regular intervals throughout inhalation, thus capturing the its fluctuation. Similar limitations are also present for cardiovascular measures, as BP and HR are highly time-variable, with possible adaptations to the respiratory challenge (Bailey et al., 2011). Continuous monitoring is advisable to improve the value of these measures as a counterpart to anxiety measures.

An important focus of this study, was the temporal variation in the TS period and testing the hypotheses that continuous self-report of pain would reveal changes during the anxiogenic inhalation, compared to control. We found that the maximal pain rating from the period, TSmax, was lower during the CO_2_ mixture compared to medical air. An increase in TES current amplitude, unsurprisingly, resulted in an increase in TSmax. However, these measures were not found to be explained by any of the anxiety measures, including aVAS, in the best mixed-effects model. This suggests that, in the early phase the pain response is less influenced or driven by an anxiety/emotional response. Importantly, this observation is distinctly different from the summative pain measurement, which shows a stronger association.

Furthermore, analysis of the rate of incline leading to TSmax, the slope of TS (TSm), confirmed an expected increase in slope with current amplitude (see Figure 4). Comparison between mixtures, revealed a reduction in the slope during the anxiogenic inhalation. As with TSmax, anxiety measures were not a predictive factor in the best mixed-effects model for TSm. Taken together, this suggests that anxiety is not a definitive driver of pain summation in the early phase of the pain report.

**Figure 4.**
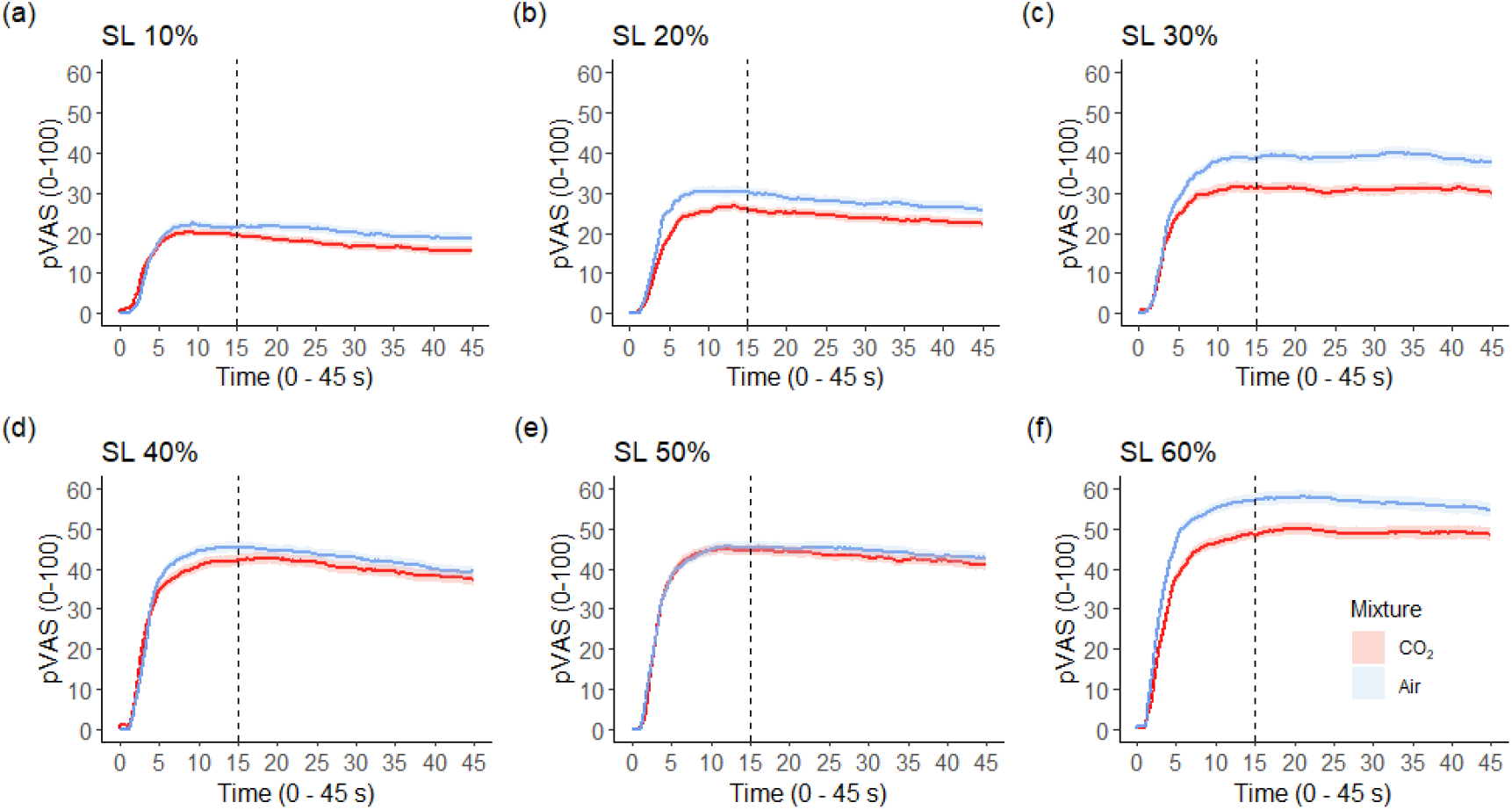
Average Pain Responses per Stimulation Level. Time in seconds is plotted on the x-axis and pain rating on visual analogue scale is plotted on the y-axis. Red line represents average response during the 7.5% CO_2_ mixture and blue line - during medical air. The grey area above and below the line represents 95% confidence intervals. SL: Stimulation level.

Given the observed association between anxiety and pain in summative measurements, this raises two possible explanations for this relationship. Firstly, it is possible that anxiety only plays a role in the interpretation of intense sensory stimuli following a sustained period, longer than the TS window. Secondly, that anxiety is perpetuated by the painful sensory stimulus, rather than being a preceding factor. The latter was tested by including pVASμ in the analyses of anxiety measures. It was found, however, that the summative pain rating did not explain any of the variance in the anxiety scores. We postulate that while a reciprocal relationship between the states of anxiety and pain exist, as suggested in similar research (Vowles et al., 2006). In our study reported pain was not universally associated with reported anxiety.

Our observations of the longer, post-TS A period, confirmed similar effects of mixture and stimulus intensity. However, individual variance was a more influential factor in the mixed effects model. This was confirmed in the analysis of the Aμ and was consistent in both Amax and Amin. Increased inter-participant variance was not unexpected, as individual variability, including age, gender, or previous experience of pain are influential of adaption to continuous pain stimulation (Cooper et al., 2013). Contrary to other continuous pain measures, analysis of Amin included aVAS in the best model. This is challenging to interpret as the range for Amin value was very large within and between participants, reflecting the variability of adaptation one may experience. We propose that the minimal rating is similar to the summative ratings in that it is sensitive to the influence of the experienced anxiety in addition to the hypercapnic inhalation. Further investigation, however, is necessary to verify that this is a reproducible effect.

While a linear relationship did not transpire between pain ratings and concurrently reported anxiety ratings, all anxiety measures, both psychological and physiological, displayed a significant increase during the anxiogenic manipulation, unlike the control inhalation of air or breaks. This increase in anxiety was independent of the TES and as mentioned above, the associated pVASμ. However, it is worth noting that the use of the CO_2_ model, and the associated screening procedure, incur a level of preselection that may influence participant characteristics. While the CO_2_ model has a lower rate of non-responders than conventional psychological methods of anxiety induction, a higher rate of drop-outs is observed. This appears to be associated with higher trait and baseline negative affectivity (Stegen et al., 1998), which may lead to premature termination of the experiment. Future work, which aims to research this relationship or would like to increase their retention rate, could use a lower concentration of CO_2_, e.g. a 5% concentration.

The central observation in this study is that the reduction in the early stage pain response, was not found to be best described by associated increase in the anxiety level. While the 7.5% CO_2_ model is a robust model of anxiety and its increase was observed here, our results indicate that CO_2_-induced anxiety is not necessarily related to reported pain. In contrast, the presence of the anxiogenic mixture alone was a definitive predictor of the reduction. While our study provides an explanation for lower pain ratings despite preserved wind-up during hypercapnia, it remains unclear what property of the inhalation triggers this effect.

In previously reported animal models, the analgesic property of the CO_2_ inhalation has been attributed to the adenosine receptors at the spinal level (Otsuguro et al., 2007). However, in human research adopting hypercapnic does not certainly establish a reduction in wind-up as measured through the spinal nociceptive reflex (Morélot-Panzini et al., 2014). The answer to why TS measures were found reduced in our study may be sourced from related literature on dyspnoea induced via inspiratory threshold loading. Such paradigm has been found to reliably inhibit the reflex (Morélot-Panzini et al., 2007). Specifically, the reduced nociceptive flexion reflex was found to be associated with both measured and self-rated respiratory stress; the reflex remained low while the respiratory challenge was described as strenuous and distressing. In our study we may not have established a linear relationship between reported anxiety and TS measures, but the anxiogenic effect evoked by the hypercapnic inhalation may still have mediated the pain measures.

## Conclusions

While temporal summation is the psychological equivalent of the physiological wind-up, it was inhibited during the inspiration of a 7.5% CO_2_ mixture. Our study reduces the gap in the anxiety literature between research on nociceptive flexion reflex and self-reported pain (Grönroos and Pertovaara, 1994).

Our data further evidence the value of continuous self-reported pain measures and suggest that they should be adopted to improve the accuracy of future research on acute experimental pain, particularly where temporal profiles of pain could be clinical applied (Staud et al., 2007).

Finally, we demonstrate important distinction between the early and late stages of acute pain experience, which suggests a more complex relationship between the concurrence of pain and anxiety. Observed reductions in pain ratings with the 7.5% CO_2_ model, which is an established model of anxiety, suggests that further research is required to disentangle the direct physiological effect of this model from the anxiety-related impact on pain.

## Acknowledgements

The authors thank the Tobacco and Alcohol Research Group, particularly Dr. Angela Attwood, from University of Bristol, UK for adhering to the principles of open science by providing technical guidance on setting up the CO_2_ model in our Faculty.

## Author Contributions

The authors have substantially contributed to the project.

