## Supplemental Material 1 for "The role of anxiety in the perception of pain: exploring the cumulative & temporal mechanisms of hypercapnic analgesia"

**Supplemental Material 1. Average Pain Measures Models**

**Table 4.** Model Statistics for Average Pain Ratings (pVASμ)

| | Frequentist Analysis | | | Bayesian Analysis | | | |
|---|---|---|---|---|---|---|---|
| | ***Coeff.*** | ***(SE)*** | **t-value** | ***Coeff.*** | ***(SD)*** | ***CI*** | |
| | | | | | | ***2.5%*** | ***97.5%*** |
| *Fixed effects* | | | | | | | |
| Intercept | 20.05 *** | (3.33) | 6.02 | 20.09 | (3.44) | 13.36 | 26.9 |
| $CO_2$ Mixture | -3.68 ** | (1.23) | 3 | -3.74 | (1.22) | -6.13 | -1.34 |
| Stimulation 20% | 6.1 ** | (1.99) | 3.06 | 6.12 | (2.03) | 2.12 | 10.08 |
| Stimulation 30% | 13.57 *** | (1.98) | 6.86 | 13.63 | (2.01) | 9.68 | 17.61 |
| Stimulation 40% | 19.88 *** | (1.96) | 10.13 | 20.01 | (2.04) | 15.95 | 24.03 |
| Stimulation 50% | 22.65 *** | (1.97) | 11.53 | 22.64 | (2) | 18.7 | 26.53 |
| Stimulation 60% | 30.40 *** | (1.99) | 15.31 | 30.38 | (2.01) | 26.39 | 34.3 |
| | $\sigma^2$ | ***(SD)*** | | $\sigma^2$ | ***(SD)*** | | |
| *Random effects* | | | | | | | |
| Participant | 180.5 | (13.43) | | 197.3 | (69.74) | 102.28 | 371.84 |
| aVAS x Participant | 55.4 | (7.44) | | 41.73 | (25.43) | .56 | 90.68 |

Note: In frequentist analysis, linear mixed model was fit by maximum likelihood, t-tests were calculated automatically using Satterthwaite approximations to degrees of freedom. In Bayesian analysis, weakly informed priors were used to estimate parameters.

Coeff./$\sigma^2$: estimated coefficient/variance; SE/SD: estimated standard error/deviation; CI is the credible interval.

*** $p < .001$, ** $p < .01$, * $p < .05$

**Table 5.** Model Statistics for Average Pain Ratings for Temporal Summation Period (TSμ)

| | Frequentist Analysis | | | Bayesian Analysis | | | |
|---|---|---|---|---|---|---|---|
| | | | | | | ***CI*** | |
| | ***Coeff.*** | ***(SE)*** | **t-value** | ***Coeff.*** | ***(SD)*** | ***2.5%*** | ***97.5%*** |
| *Fixed effects* | | | | | | | |
| Intercept | 17.99 *** | (2.63) | 6.84 | 17.99 | (2.73) | 12.59 | 23.36 |
| $CO_2$ Mixture | -2.9 ** | (1.07) | 2.72 | -2.91 | (1.04) | -4.92 | -.85 |
| Stimulation 20% | 4.72 ** | (1.7) | 2.79 | 4.72 | (1.71) | 1.35 | 8.08 |
| Stimulation 30% | 8.97 *** | (1.67) | 5.37 | 9.01 | (1.7) | 5.69 | 12.36 |
| Stimulation 40% | 15.5 *** | (1.65) | 9.42 | 15.59 | (1.71) | 12.21 | 18.96 |
| Stimulation 50% | 18.16 *** | (1.65) | 10 | 18.16 | (1.69) | 14.91 | 21.5 |
| Stimulation 60% | 23.19 *** | (1.68) | 13.8 | 23.18 | (1.69) | 19.86 | 26.52 |
| | $\sigma^2$ | ***(SD)*** | | $\sigma^2$ | ***(SD)*** | | |
| *Random effects* | | | | | | | |
| Participant | 106.39 | (10.31) | | 115.68 | (41.47) | 60 | 218.44 |
| aVAS x Participant | 64.06 | (8) | | 57.65 | (18.65) | 16.52 | 91.16 |

Note: In frequentist analysis, linear mixed model was fit by maximum likelihood, t-tests were calculated automatically using Satterthwaite approximations to degrees of freedom. In Bayesian analysis, weakly informed priors were used to estimate parameters.

Coeff./$\sigma^2$: estimated coefficient/variance; SE/SD: estimated standard error/deviation; CI is the credible interval.

*** $p < .001$, ** $p < .01$, * $p < .05$

**Table 6**. Model Statistics for Average Pain Ratings for Adaptation Period (Aμ)

| | Frequentist Analysis | | | Bayesian Analysis | | | |
|---|---|---|---|---|---|---|---|
| | | | | | | *CI* | |
| | ***Coeff.*** | ***(SE)*** | **t-value** | ***Coeff.*** | ***(SD)*** | ***2.5%*** | ***97.5%*** |
| *Fixed effects* | | | | | | | |
| Intercept | 21.09 *** | (3.79) | 5.56 | 21.06 | (3.94) | 13.36 | 28.84 |
| $CO_2$ Mixture | -4.11 ** | (1.36) | 3.02 | -4.18 | (1.34) | -6.85 | -1.56 |
| Stimulation 20% | 6.81 ** | (2.24) | 3.04 | 6.84 | (2.23) | 2.54 | 11.27 |
| Stimulation 30% | 15.87 *** | (2.23) | 7.13 | 15.89 | (2.25) | 11.48 | 20.36 |
| Stimulation 40% | 22.05 *** | (2.21) | 9.97 | 22.15 | (2.28) | 17.67 | 26.6 |
| Stimulation 50% | 24.85 *** | (2.21) | 11.22 | 24.83 | (2.23) | 20.52 | 29.19 |
| Stimulation 60% | 34.01 *** | (2.23) | 15.24 | 33.98 | (2.24) | 29.51 | 38.38 |
| | $\sigma^2$ | ***(SD)*** | | $\sigma^2$ | ***(SD)*** | | |
| *Random effects* | | | | | | | |
| Participant | 236.63 | (15.38) | | 257.11 | (91.19) | 132.29 | 487.48 |
| aVAS x Participant | 51.64 | (7.19) | | 38.55 | (27.85) | .14 | 96.34 |

Note: In frequentist analysis, linear mixed model was fit by maximum likelihood, t-tests were calculated automatically using Satterthwaite approximations to degrees of freedom. In Bayesian analysis, weakly informed priors were used to estimate parameters.

Coeff./$\sigma^2$: estimated coefficient/variance; SE/SD: estimated standard error/deviation; CI is the credible interval.

*** $p < .001$, ** $p < .01$, * $p < .05$
