## Supplemental Material 2 for "The role of anxiety in the perception of pain: exploring the cumulative & temporal mechanisms of hypercapnic analgesia"

**Supporting Material 2. Continuous Pain Measures Models**

**Table 7.** Model Statistics for Maximal Pain Rating during Temporal Summation (TSmax)

| | Frequentist Analysis | | | Bayesian Analysis | | | |
|---|---|---|---|---|---|---|---|
| | | | | | | CI | |
| | *Coeff.* | *(SE)* | t-value | *Coeff.* | *(SD)* | *2.5%* | *97.5%* |
| *Fixed effects* | | | | | | | |
| Intercept | 26.49 *** | (3.59) | 7.37 | 26.36 | (3.89) | 18.57 | 33.92 |
| $CO_2$ Mixture | -4.56 *** | (1.21) | 3.78 | -4.56 | 1.21 | -6.94 | -2.13 |
| Stimulation 20% | 7.1 *** | (2.09) | 3.39 | 7.03 | (2.09) | 3 | 11.16 |
| Stimulation 30% | 13.66 *** | (2.09) | 6.53 | 13.6 | (2.1) | 9.48 | 17.76 |
| Stimulation 40% | 21.53 *** | (2.09) | 10.29 | 21.44 | (2.12) | 17.38 | 25.6 |
| Stimulation 50% | 23.16 *** | (2.09) | 11.07 | 23.08 | (2.13) | 18.97 | 27.35 |
| Stimulation 60% | 30.38 *** | (2.09) | 14.52 | 30.29 | (2.15) | 26.06 | 34.44 |
| | $\sigma^2$ | *(SD)* | | $\sigma^2$ | *(SD)* | | |
| *Random effects* | | | | | | | |
| Participant | 217.4 | (14.74) | | 253.22 | (91.06) | 133.63 | 480.48 |

Note: In frequentist analysis, linear mixed model was fit by maximum likelihood, t-tests were calculated automatically using Satterthwaite approximations to degrees of freedom. In Bayesian analysis, weakly informed priors were used to estimate parameters.

Coeff./$\sigma^2$: estimated coefficient/variance; SE/SD: estimated standard error/deviation; CI is the credible interval.

*** $p < .001$, ** $p < .01$, * $p < .05$

**Table 8.** Model Statistics for Average Slope during Temporal Summation (TSm)

| | Frequentist Analysis | | | Bayesian Analysis | | | |
|---|---|---|---|---|---|---|---|
| | ***Coeff.*** | ***(SE)*** | **t-value** | ***Coeff.*** | ***(SD)*** | ***CI*** ***2.5%*** | ***97.5%*** |
| *Fixed effects* | | | | | | | |
| Intercept | 5.2 *** | (.8) | 6.46 | 6.58 | (.83) | 4.94 | 8.21 |
| $CO_2$ Mixture | -1.37 ** | (.5) | 2.71 | -1.38 | (.51) | -2.36 | -.38 |
| Stimulation 20% | .22 | (.88) | .25 | .19 | (.86) | -1.48 | 1.87 |
| Stimulation 30% | .98 | (.88) | 1.11 | .95 | (.88) | -.79 | 2.65 |
| Stimulation 40% | 2.16 * | (.88) | 2.46 | 2.14 | (.87) | .41 | 3.84 |
| Stimulation 50% | 2.13 * | (.88) | 2.42 | 2.11 | (.88) | .39 | 3.8 |
| Stimulation 60% | 1.92 * | (.88) | 2.18 | 1.88 | (.88) | .16 | 3.59 |
| | $\sigma^2$ | ***(SD)*** | | $\sigma^2$ | ***(SD)*** | | |
| *Random effects* | | | | | | | |
| Participant | 6.49 | (2.55) | | 4.78 | (2.15) | 1.87 | 10.22 |

Note: In frequentist analysis, linear mixed model was fit by maximum likelihood, t-tests were calculated automatically using Satterthwaite approximations to degrees of freedom. In Bayesian analysis, weakly informed priors were used to estimate parameters.

Coeff./$\sigma^2$: estimated coefficient/variance; SE/SD: estimated standard error/deviation; CI is the credible interval.

*** $p < .001$, ** $p < .01$, * $p < .05$

**Table 9.** Model Statistics for Maximal Pain Ratings during Adaptation Period (Amax)

| | Frequentist Analysis | | | Bayesian Analysis | | | |
|---|---|---|---|---|---|---|---|
| | | | | | | *CI* | |
| | ***Coeff.*** | ***(SE)*** | **t-value** | ***Coeff.*** | ***(SD)*** | ***2.5%*** | ***97.5%*** |
| *Fixed effects* | | | | | | | |
| Intercept | 24.88 *** | (3.75) | 6.64 | 24.86 | (3.99) | 16.95 | 32.47 |
| $CO_2$ Mixture | -3.92 ** | (1.28) | 3.06 | -3.92 | (1.29) | -6.5 | -1.41 |
| Stimulation 20% | 7.81 *** | (2.22) | 3.52 | 7.69 | (2.23) | 3.37 | 12.04 |
| Stimulation 30% | 17.15 *** | (2.22) | 7.73 | 17.04 | (2.24) | 12.72 | 21.45 |
| Stimulation 40% | 23.76 *** | (2.22) | 10.71 | 23.66 | (2.2) | 19.46 | 27.97 |
| Stimulation 50% | 25.21 *** | (2.22) | 11.36 | 25.09 | (2.21) | 20.76 | 29.38 |
| Stimulation 60% | 34.46 *** | (2.22) | 15.53 | 34.32 | (2.2) | 30 | 38.65 |
| | $\sigma^2$ | ***(SD)*** | | $\sigma^2$ | ***(SD)*** | | |
| *Random effects* | | | | | | | |
| Participant | 234.6 | (15.32) | | 268.36 | (91.89) | 139.39 | 490.32 |

Note: In frequentist analysis, linear mixed model was fit by maximum likelihood, t-tests were calculated automatically using Satterthwaite approximations to degrees of freedom. In Bayesian analysis, weakly informed priors were used to estimate parameters.

Coeff./$\sigma^2$: estimated coefficient/variance; SE/SD: estimated standard error/deviation; CI is the credible interval.

*** $p < .001$, ** $p < .01$, * $p < .05$

**Table 10.** Model Statistics for Minimal Pain Ratings during Adaptation Period (Amin)

| | Frequentist Analysis | | | Bayesian Analysis | | | |
|---|---|---|---|---|---|---|---|
| | | | | | | *CI* | |
| | ***Coeff.*** | ***(SE)*** | **t-value** | ***Coeff.*** | ***(SD)*** | ***2.5%*** | ***97.5%*** |
| *Fixed effects* | | | | | | | |
| Intercept | 21.09 *** | (3.79) | 5.56 | 21.06 | (3.94) | 13.36 | 28.84 |
| $CO_2$ Mixture | -4.11 ** | (1.36) | 3.02 | -4.18 | (1.34) | -6.85 | -1.56 |
| Stimulation 20% | 6.81 ** | (2.24) | 3.04 | 6.84 | (2.23) | 2.54 | 11.27 |
| Stimulation 30% | 15.87 *** | (2.23) | 7.13 | 15.89 | (2.25) | 11.48 | 20.36 |
| Stimulation 40% | 22.05 *** | (2.21) | 9.97 | 22.15 | (2.28) | 17.67 | 26.6 |
| Stimulation 50% | 24.85 *** | (2.21) | 11.22 | 24.83 | (2.23) | 20.52 | 29.19 |
| Stimulation 60% | 34.01 *** | (2.23) | 15.24 | 33.98 | (2.24) | 29.51 | 38.38 |
| | $\sigma^2$ | ***(SD)*** | | $\sigma^2$ | ***(SD)*** | | |
| *Random effects* | | | | | | | |
| Participant | 236.63 | (15.38) | | 257.11 | 91.19 | 132.29 | 487.48 |
| aVAS x Participant | 51.64 | (7.19) | | 38.55 | (27.85) | .14 | 96.34 |

Note: In frequentist analysis, linear mixed model was fit by maximum likelihood, t-tests were calculated automatically using Satterthwaite approximations to degrees of freedom. In Bayesian analysis, weakly informed priors were used to estimate parameters.

Coeff./$\sigma^2$: estimated coefficient/variance; SE/SD: estimated standard error/deviation; CI is the credible interval.

*** $p < .001$, ** $p < .01$, * $p < .05$
